# Linking genotypes with multiple phenotypes in single-cell CRISPR screens

**DOI:** 10.1101/658146

**Authors:** Lin Yang, Yuqing Zhu, Hua Yu, Sitong Chen, Yulan Chu, He Huang, Jin Zhang, Wei Li

**Affiliations:** Center for Genetic Medicine Research, Children’s National Medical Center, 111 Michigan Ave NW, Washington, DC 20010; Department of Genomics and Precision Medicine, George Washington University, 111 Michigan Ave NW, Washington, DC 20010; Department of Biochemistry & Molecular Medicine, George Washington University, 2300 Eye St., NW, Washington, D.C. 20037; Center for Stem Cell and Regenerative Medicine, Department of Basic Medical Sciences, The First Affiliated Hospital, College of Medicine, Zhejiang University, Hangzhou, Zhejiang 310058, China; Institute of Hematology, Zhejiang University, Hangzhou, Zhejiang 310058, China

**Author notes:** Equal contributions (Lin Yang, Yuqing Zhu, Hua Yu).

## Abstract

CRISPR/Cas9 based functional screening coupled with single-cell RNA-seq (“single-cell CRISPR screening”) unravels gene regulatory networks and enhancer-gene regulations in a large scale. We propose scMAGeCK, a computational framework to systematically identify genes and non-coding elements associated with multiple expression-based phenotypes in single-cell CRISPR screening. scMAGeCK identified genes and enhancers that modulate the expression of a known proliferation marker, MKI67 (Ki-67), a result that resembles the outcome of proliferation-linked CRISPR screening. We further performed single-cell CRISPR screening on mouse embryonic stem cells (mESC), and identified key genes associated with different pluripotency states. scMAGeCK enabled an unbiased construction of genotype-phenotype network, where multiple phenotypes can be regulated by different gene perturbations. Finally, we studied key factors that improve the statistical power of single-cell CRISPR screens, including target gene expression and the number of guide RNAs (gRNAs) per cell. Collectively, scMAGeCK is a novel and effective computational tool to study genotype-phenotype relationships at a single-cell level.

## Introduction

Pooled genetic screens based on CRISPR/Cas9 genome engineering system is a widely used method to study the functions of thousands of genes or non-coding elements in one single experiment (Shalem et al., 2014; Wang et al., 2014; Zhou et al., 2014). Recent CRISPR screening combined with single-cell RNA-seq (scRNA-seq) provides a powerful method to monitor gene expression changes in response to perturbation at a single-cell level. These technologies, including Perturb-seq (Adamson et al., 2016; Dixit et al., 2016), CRISP-seq (Jaitin et al., 2016), Mosaic-seq (Xie et al., 2017), CROP-seq (Datlinger et al., 2017), etc, enabled a large-scale investigation of gene regulatory networks, genetic interactions and enhancer-gene regulations in one experiment.

CRISPR screening coupled with scRNA-seq, which will be referred to as “single-cell CRISPR screening”, enables detecting the expression changes of whole transcriptome at a single-cell level. One can potentially search for perturbed genomic elements that lead to the differential expression of certain gene of interest. This approach resembles a fluorescence-activated cell sorting (FACS) experiment, where single cells are separated into groups of high (or low) expression of certain marker. Such “virtual FACS” experiment (Xie et al., 2017) can be performed on unlimited numbers of phenotypes, represented by the expressions of genes (or gene signatures). Therefore, single-cell CRISPR screening greatly increases the limitation of traditional screening experiment, where only one phenotype can be tested. However, few efforts are made to evaluate this approach, and no computational methods are available for the “virtual FACS” analysis based on single-cell CRISPR screening data.

Here we present scMAGeCK, a computational framework to systematically identify genes (and non-coding elements) associated with multiple phenotypes in single-cell CRISPR screening data. scMAGeCK is based on our previous MAGeCK models for pooled CRISPR screens (Li et al., 2014; 2015; Wang et al., 2019), but further extends to scRNA-seq as the readout of the screening experiment. scMAGeCK consists of two modules: scMAGeCK-Robust Rank Aggregation (RRA), a sensitive and precise algorithm to detect genes whose perturbation links to one single marker expression; and scMAGeCK-LR, a linear-regression based approach that unravels the regulatory relationship of thousands of gene expressions from cells with multiple perturbations.

We demonstrated the ability of scMAGeCK to perform functional analysis from single-cell CRISPR screens. We applied scMAGeCK on public datasets generated from CROP-seq (Datlinger et al., 2017), a widely used protocol for single-cell CRISPR screening (Hill et al., 2018; Shifrut et al., 2018; Gasperini et al., 2019). Gene expression clustering is a typical approach to evaluate the effects of gene perturbations. We found that for all the datasets, only one to two genes are enriched in clusters, while scMAGeCK identified 25%-95% targets whose expressions are down-regulated upon knockout with statistical significance. Applying this approach to phenotypes, we identified oncogenic and tumor-suppressor genes and enhancers, by simply testing their associations with MKI67 (Ki-67), a commonly used proliferation marker. The output of scMAGeCK further enabled the construction of a complex genotype-phenotype network. We tested our scMAGeCK approach on mouse embryonic stem cells (mESC), and identified key genes associated with different pluripotency states.

Finally, we studied key factors that determine the statistical power of single-cell CRISPR screens. The efficiency of gene knockouts (or knockdowns) varies between different targets and different single cells. Highly expressed target genes tend to have a stronger effect of down-regulation compared with moderately or lowly expressed targets. Screens with high multiplicity of infection (MOI), where multiple sgRNAs enter into one cell, have improved sensitivity and specificity compared with screens performed in low MOI.

## Results

### scMAGeCK method overview

We previously developed MAGeCK and MAGeCK-VISPR, two algorithms to model gene knockouts from genome-wide CRISPR/Cas9 screens (Li et al., 2014; 2015). MAGeCK models the read counts of single guide RNAs (sgRNAs) using a Negative Binomial (NB) distribution, and prioritizes genes with a revised robust rank aggregation algorithm (alpha-RRA, (Kolde et al., 2012)). The alpha parameter introduced in MAGeCK is used to determine significant and non-significant gRNAs. In addition, “MAGeCK-VISPR” models complex experimental designs using a generalized linear model, and uses an Expectation-Maximization (EM) approach to optimize all the parameters.

scMAGeCK applies the statistical models of MAGeCK and MAGeCK-VISPR to single-cell CRISPR screening data. scMAGeCK includes two modules, scMAGeCK-RRA and scMAGeCK-LR (Fig. 1a). To identify genes whose perturbation associated with the expression of a gene of interest, scMAGeCK-RRA first ranks single cells according to the target gene expression. Next, scMAGeCK uses RRA to test whether single cells with particular gene perturbation are enriched in a higher (or lower) expression of the target. The alpha parameter is set to exclude single cells whose marker expression is zero, possibly due to dropout events. Another module, scMAGeCK-LR, simultaneously investigates the effects of all possible gene expressions. scMAGeCK-LR uses a linear regression model to calculate the “selection” score, similar to “log-fold change”, that describes the degree of perturbations (see Methods for more details).

**Figure 1.**
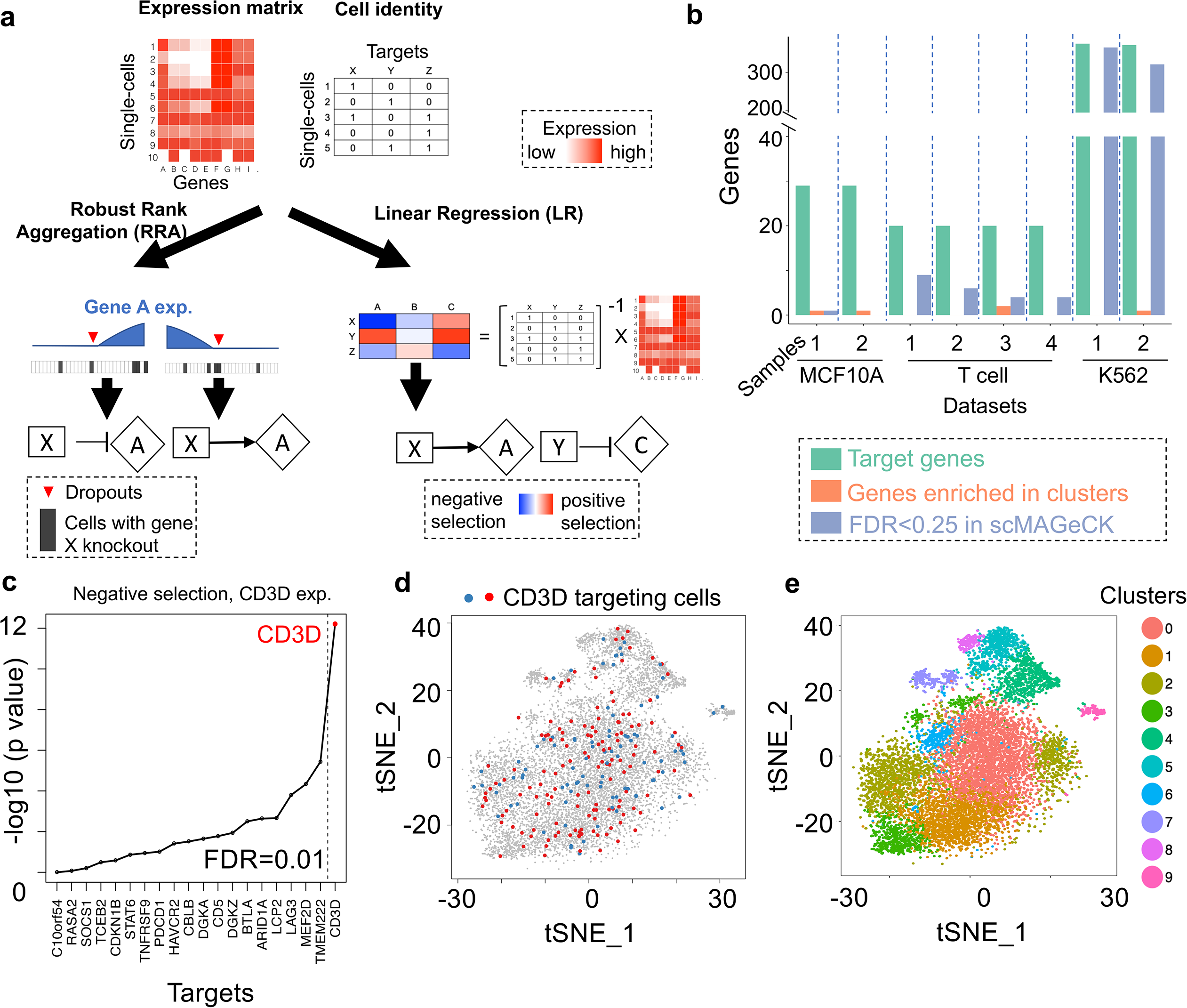
scMAGeCK pipeline and a comparison with clustering analysis on single-cell CRISPR screens. **a,** An overview of the scMAGeCK pipeline. The input of scMAGeCK includes a scaled expression matrix of all genes in all single-cells, together with cell identity information on the targets of each single cell. scMAGeCK includes two modules: RRA and LR. RRA infers gene regulatory relationship on certain gene expression (e.g., gene A) using the rankings of single cells, and takes dropout events into consideration. LR infers the gene regulatory network on all possible gene expressions. **b,** A comparison of scMAGeCK with clustering analysis on three different public CROP-seq datasets. The total number of target genes, genes that are enriched in certain cluster, and genes whose down-regulation is considered as statistically significant (FDR<0.25) are shown. Gene A is considered enriched in certain cluster are defined as single cells carrying gene A knockout consists of >20% total cells in that cluster. **c,** The ranking of genes in reducing CD3D expression in the T cell CROP-seq dataset. **d-e,** Single-cells carrying CD3D sgRNA (d), and the clustering of all single cells in the T cell CROPseq dataset (e), using t-Distributed Stochastic Neighbor Embedding (t-SNE) analysis.

scMAGeCK-RRA and scMAGeCK-LR provides two different approaches for single-cell CRISPR screening data. scMAGeCK-RRA is a sensitive approach to detect subtle and nonlinear expression changes, while scMAGeCK-RRA simultaneously tests the expressions of thousands of genes, and is able to deal with cells targeted by multiple sgRNAs.

### Comparisons with clustering analysis

A typical approach to analyze perturbation effect in single-cell CRISPR screening is “enrichment by clustering”: users first cluster single cells based on their gene expression patterns, then check whether certain sgRNAs are enriched in one or more of these clusters. We applied this approach to several public CROP-seq datasets performed on different cell types, including breast epithelial cells (MCF10A), un-stimulated and stimulated primary human T cells, and myelogenous leukemia cells (K562) (Hill et al., 2018; Shifrut et al., 2018; Gasperini et al., 2019). The number of perturbed genes or enhancers vary from around 20 (MCF10A and T cell) to over one thousand (K562). We found that the clustering approach only identified 1-2 genes whose sgRNAs are enriched in certain clusters (Fig. 1b). The small number of enriched targets in clusters thus limits downstream analysis, including the evaluation of knockout efficiency.

Instead of clustering analysis, we used scMAGeCK-RRA to investigate whether target gene knockout reduces their expressions. In two out of three datasets, we found that 25%-95% of the target genes have reduced expressions with statistical significance, a demonstration that scMAGeCK-RRA better captures the effect of gene perturbation than the clustering analysis. For example, CD3D knockout strongly reduces CD3D expressions in single cells (Fig. 1c), while cells targeting CD3D are not enriched in any clusters (Fig. 1d-e).

### Identification of known oncogenic and tumor suppressor genes and enhancers

We first used scMAGeCK-RRA to identify genes that modulate the expression of Ki-67 (*MKI67*), a commonly used marker for cell proliferation. In MCF10A CROP-seq, the knockout of *TP53* tumor suppressor gene strongly induced MKI67 expression in corresponding single cells (adjusted p value=1.5e-4), Other gene knockouts (*RUNX1, CDH1* and *ARID1B*) have similar effect, consistent with their reported tumor suppressor roles in breast cancer or other cancer types (Pećina-Slaus, 2003; Khursheed et al., 2013; Hong et al., 2018) (Fig. 2a). On the other hand, four gene knockouts significantly reduce Ki-67 expression (Fig. 2b). Among those, CHEK1 is a checkpoint kinase that is essential for normal and cancer cells (Zhang and Hunter, 2014), GATA3 is a critical transcription factor with known oncogenic role (Mehra et al., 2005), and RAD51 has been reported as an oncogene with elevated expression in multiple cancer types including breast cancer (Maacke et al., 2000). CASP8 has multiple functions in different contexts (Stupack, 2013), with a possible essential role in breast cancer cell lines (De Blasio et al., 2016).

**Figure 2.**
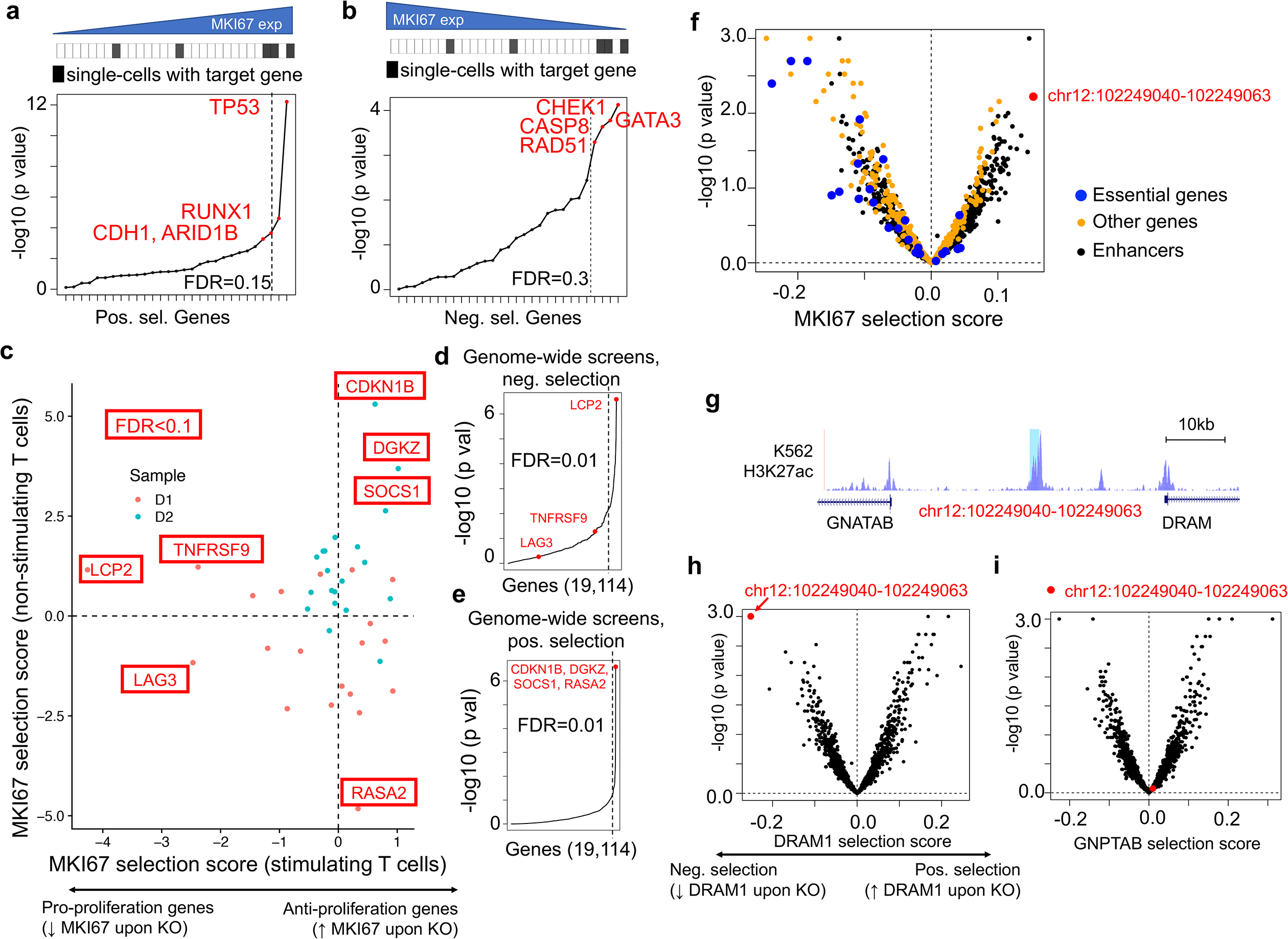
Associations with the expression of Ki-67 (MKI67), a proliferation marker. **a-b,** The rankings of genes that are positively (a) or negatively selected (b) on MKI67 expression. Here, positive selection in (a) indicates single cells with certain target gene knockout (black rectangle on the top) have higher MKI67 expression. **c,** The MKI67 selection score in stimulating and non-stimulating T cells in the T cell CROP-seq dataset. Genes performed in two different patient samples (D1/D2) are marked with different colors, and genes with FDR<0.1 are highlighted. The selection score is calculated based on the p values reported from RRA, with direction depending on whether it is a positively (or negatively) selected genes. See Methods for more details. **d-e,** The ranking of selected genes in c in genome-wide CRISPR screening, including negative selection ranking (d) and positive selection ranking (e). **f,** The MKI67 selection score and p values calculated from scMAGeCK-LR in K562 dataset. Essential genes (ribosomal subunits and proteasomes) are marked in blue, while the tumor-suppressor-like enhancer of interest (chr12:102249040-102249063) is highlighted in red. **g,** The chromosome view of the enhancer chr12:102249040-102249063. h-i, the DRAM1 and GNPTAB selection score and their corresponding p values. The enhancer chr12:102249040-102249063 is highlighted in red.

In the T cell CROP-seq dataset, we identified different genes that regulate *MKI67* expression in non-stimulating and stimulating T cells (Fig. 2c), and compared their roles in genome-wide CRISPR screens (Fig. 2d-e). Here, we defined a “selection score” based on the p values calculated by scMAGeCK-RRA to describe the direction (and the degree) of *MKI67* regulation (see Methods for more details). Among those, four genes play anti-proliferation roles in stimulating T cells (*CDKN1B, DGKZ, SOCS1* and *RASA2*). All these genes are top positively selected hits in genome-wide CRISPR screens (Fig. 2e). LCP2, the strongest negative selection hit in genome-wide screens (Fig. 2d), is also identified as the top pro-proliferation gene, consistent with its essential role in T cell function (Shifrut et al., 2018). TNFRSF9 (CD137) is a co-stimulatory factor in T cells whose knockout reduces *MKI67* expression but is not identified in genome-wide CRISPR screens (Fig. 2d). In contrast, LAG3, an immune checkpoint receptor, paradoxically reduces MKI67 expression upon knockout, a demonstration that different platforms may provide different results.

We next studied the expression of Ki-67 in the K562 CROP-seq dataset, where each cell is targeted by an average of twenty sgRNAs (Gasperini et al., 2019). scMAGeCK-LR is used for the analysis, as scMAGeCK-RRA is not suitable for cells targeted by multiple gRNAs. Overall, knocking down essential genes, including ribosomal subunits, proteasomes, *etc*., reduced MKI67 expression (Fig. 2f), consistent with their critical roles in cell functions. Several enhancers are among the top candidates whose perturbation changed Ki-67 expression (Fig. 2f). Among those, chr12:102249040-102249063 a putative enhancer that negatively regulate Ki-67 expression. This enhancer is located in the intergenic region of chromosome 12 with strong H3K27ac signals, proximal to the transcription start site (TSS) of two protein coding genes (*GNPTAB* and *DRAM1*, Fig. 2g). To further identify the target genes, we ranked all genes/enhancers based on their perturbation effects on *GNPTAB* and *DRAM1* expressions (Fig. 2h-i). chr12:102249040-102249063 is among the top hits on reducing the expression of *DRAM1* (but not *GNPTAB*). Indeed, *DRAM1* (DNA damage regulated autophagy modulator 1) is a tumor suppressor gene with decreased expression in various tumors, and is required for the induction of autophagy by the p53 pathway (Crighton et al., 2006). Collectively, these results demonstrated that oncogenic and tumor suppressor genes (and enhancers) can be readily identified by testing their associations with Ki-67 using scMAGeCK.

### Investigating multiple phenotypes using scMAGeCK

We set out to use scMAGeCK to study multiple phenotypes beyond proliferation. In MCF10A CROP-seq dataset, we studied the effect of gene knockouts on apoptosis, as doxorubicin is known to induce apoptosis in normal and tumor cells (Wang et al., 2004). We used the average expression of genes in an apoptosis signature in the MSigDB database (Subramanian et al., 2005) as the readout. These genes are down-regulated in a breast cancer cell line (MEA) undergoing apoptosis in response to doxorubicin (Graessmann et al., 2007), a system mostly resemble the experimental conditions in MCF10A CROP-seq. Under the false discovery rate 0.1 cutoff, we found three genes that significantly modulate the expressions of apoptosis signatures in two conditions (doxorubicin treatment or mock treatment, Fig. 3a). Among those, *TP53* consistently served as a pro-apoptosis gene, consistent with its critical role in apoptosis. Interestingly, BRCA1 serves as an anti-apoptosis gene in the normal MCF10A cells, consistent with previous reports that BRCA1 loss triggers apoptosis and BRCA1 deletion causes growth inhibition in MCF10A (Deng and Wang, 2003; You et al., 2004).

**Figure 3.**
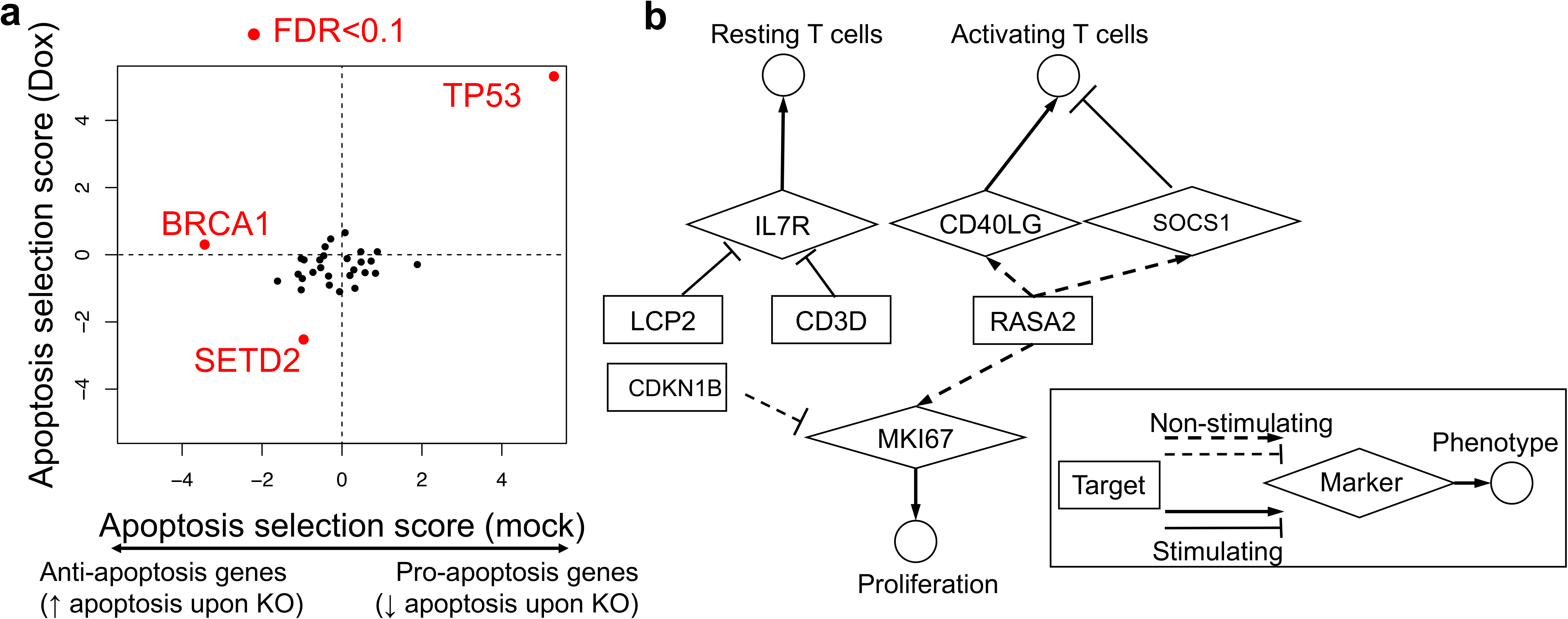
scMAGeCK on other phenotypes. **a,** The apoptosis selection score of different genes in MCF10A CROP-seq datasets treated with doxorubicin (y axis) and with mock control (x axis). Genes with FDR<0.1 are highlighted in red. Here the average expression of signature genes in an apoptosis gene set in MSigDB is served as marker. The signature comes from genes that are down-regulated in a breast cancer cell line (ME-A) undergoing apoptosis in response to doxorubicin (ID: GRAESSMANN_APOPTOSIS_BY_DOXORUBICIN_DN). **b,** The target-marker-phenotype network in T cell CROP-seq dataset. Target genes are genes that are screened in CROP-seq dataset, while markers that are known to be associated with resting T cells, activating T cells and proliferation are selected. Gene regulatory relationship from non-stimulating and stimulated cells are shown in dashed and solid lines, respectively.

In T cell CROP-seq data, we chose markers that are known to associate multiple phenotypes in T cell functions, including resting (IL7R), activating T cells (CD40LG, SOCS1), and proliferation (MKI67). The outputs of scMAGeCK enabled an unbiased construction of genotype-phenotype network in non-stimulating and stimulating T cells (Fig. 3b). Among these, LCP2 and CD3D knockout significantly increases IL7R, consistent with their essential roles in T cell stimulation. Some genes may have opposite roles in different conditions; for example, RASA2 is a positive regulator of MKI67 in non-stimulating cells (Fig. 3b) but a negative regulator in stimulating cells and genome-wide screens (Fig. 2 c-e). This genotype-phenotype network provides an intuitive approach to study gene functions in different contexts.

### scMAGeCK identified key genes associated with different pluripotency states of embryonic stem cells

Having demonstrated the ability of scMAGeCK to perform functional analysis of multiple phenotypes, we performed CROP-seq experiments to interrogate genes that are critical for mouse embryonic stem cell (mESC) pluripotency and differentiation. The pluripotent state of the mESC is highly dynamic, including a more primitive naïve state and a primed state ready for differentiation (Hackett and Surani, 2014). As they represent two key different developmental stages of pre and post-implantation embryos, it is important to understand what factors regulate these two states. We thus designed 45 guides to perturb fifteen genes including naive and primed pluripotency-associated transcription factors and metabolic genes. CROP-seq experiments were performed with samples in the two states of mESC (naïve and primed), respectively (see Methods). Overall, we obtained the transcriptome profiles of ~2,000 cells per sample using the InDrop platform (Klein et al., 2015). t-distributed stochastic neighbor embedding (t-SNE) clustering demonstrated a clear separation of both states, not batches (Fig. 4a). Known markers are selectively expressed in each state, including *Nanog* in the naïve state, and *Dnmt3b* in the primed state (Fig. 4b), respectively.

**Figure 4.**
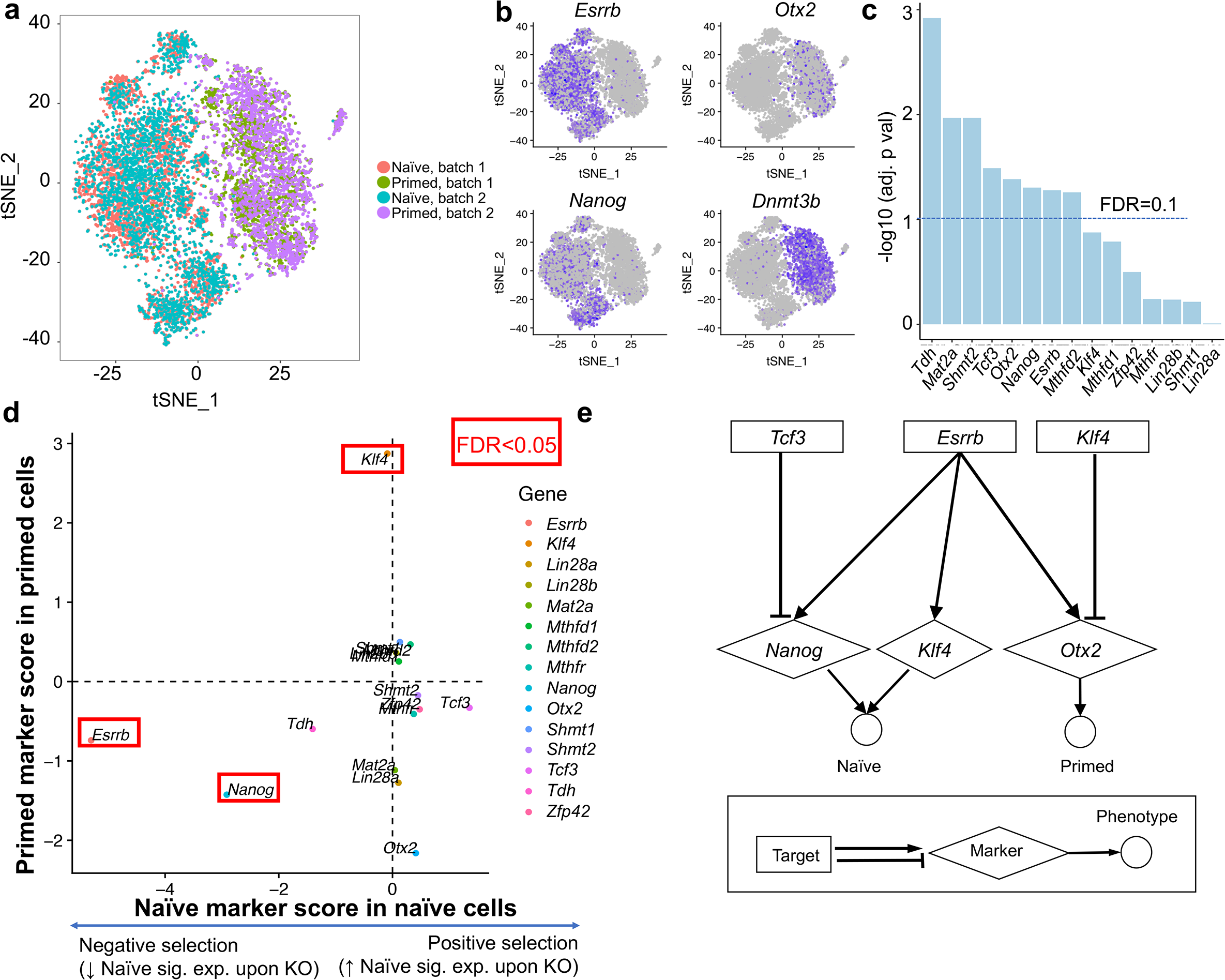
CROP-seq on mouse embryonic stem cells (mESC) uncovered known regulators for stem cell differentiation. **a,** The t-SNE plot on single-cell expression profiles in naïve or primed states in two batches. **b,** Selected marker expression, including *Esrrb, Nanog* (naïve markers), *Otx2* and *Dnmt3b* (primed markers). **c,** The adjusted p values, calculated from scMAGeCK-RRA, on target gene down-regulation. **d,** The naïve marker scores as well as primed marker scores in their corresponding cell states. Genes with FDR<0.05 are highlighted in red. Naïve marker score is based on the average expression of four naïve marker genes: *Nanog, Esrrb, Klf4* and *Tdh*, while primed marker score is based on *Otx2* expression. **e,** The target-marker-phenotype network constructed from scMAGeCK results.

Consistent with the results from public CROP-seq datasets, clustering analysis only identified 2 out of 15 target genes that are enriched in certain clusters (Supplementary Fig. 1). In contrast, scMAGeCK-RRA identified 8 out of 15 genes whose expression is reduced upon knockout with statistical significance (Fig. 4c).

We next investigated the effect of individual gene knockout on both states, using the expression of known naïve and primed markers. To this end, we used the expression of *Otx2*, a primed state-specific gene (Buecker et al., 2014) and a combined expression of *Nanog, Esrrb, Klf4* and *Tdh*, four naïve markers, as the readout (Dunn et al., 2014). The scores of both markers are shown for naïve and primed cells, respectively (Fig. 4d). Among those, *Nanog* knockout significantly reduced the naïve marker expression, consistent with its critical role in naïve pluripotency (Chambers et al., 2007). *Esrrb* knockout decreases, whereas *Tcf3* knockout increases the naive marker expressions, consistent with the previous report that *Tcf3* inhibits naïve state through *Esrrb* (Martello et al., 2012). In the primed state sample, *Klf4* knockout increases primed markers, demonstrating its role in maintaining naive state and preventing differentiation (Zhang et al., 2010).

Based on the known functions of perturbed genes, we built a target-marker-phenotype network that describes the gene regulatory network in both cell types (Fig. 4e). This network is consistence with previously reported naive and primed regulatory network (Weinberger et al., 2016).

### High target expression and high MOI improves the power of single-cell CRISPR screening

We set out to determine factors that affect the statistical power of single-cell CRISPR screening. We first determine whether the expression of target gene is reduced in corresponding single cells, an indication of target knockout efficiency. Different levels of down-regulation are observed in different datasets and samples (Fig. 1b). Overall, we observed a strong correlation between the effect of down-regulation (measured by the negative selection p values from scMAGeCK) and median gene expression in all datasets (Fig. 5a-b, Supplementary Fig. 2). Genes that are highly expressed are more likely to have a strong down-regulation. For example, in mESC CROP-seq dataset, targets may undergo different down-regulation effect in different states (Fig. 5b). *Tdh*, a highly expressed gene in naïve but not in primed cells, demonstrates strong down-regulation effect only in the naïve state (Fig. 5c-d).

**Figure 5.**
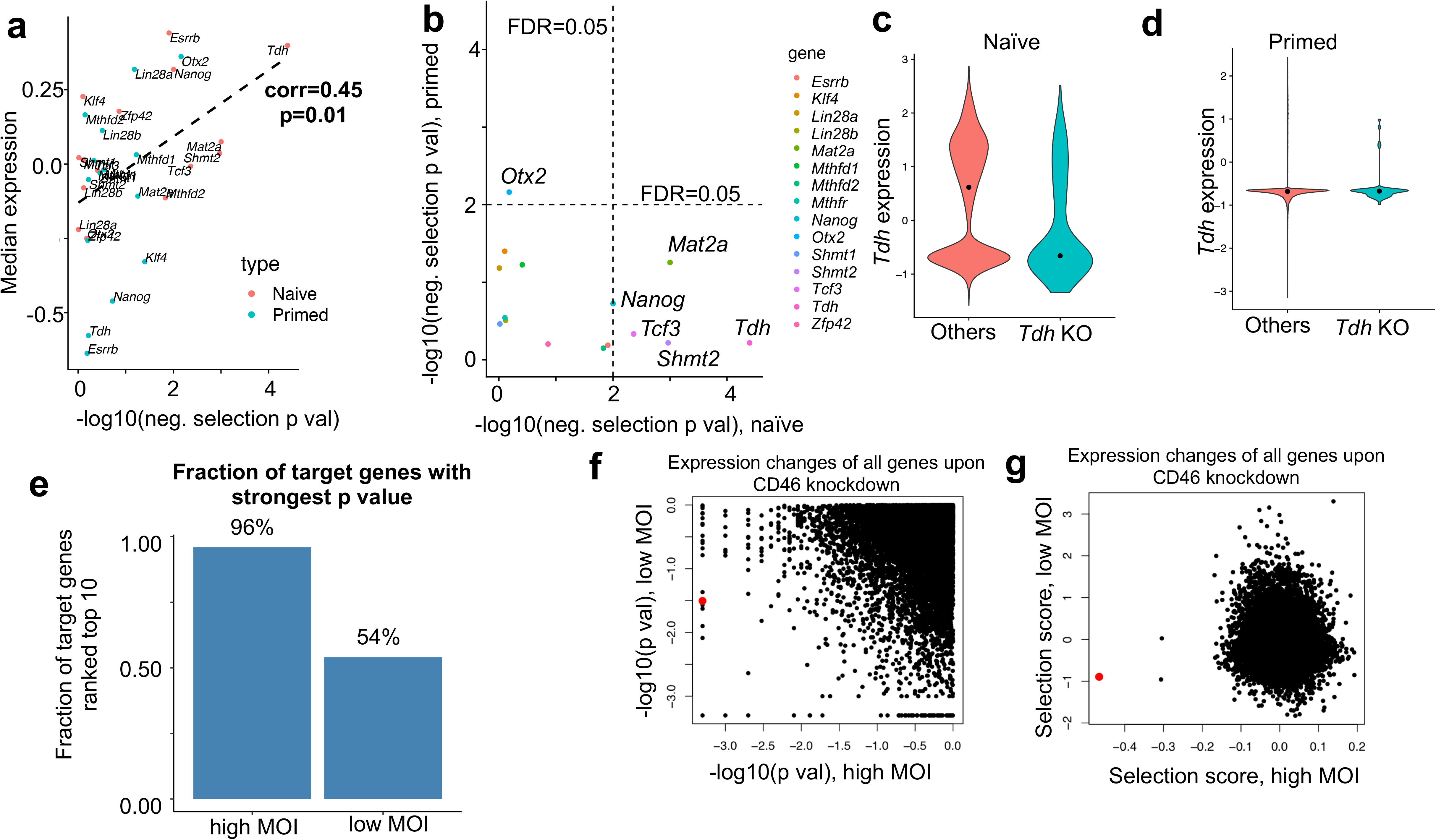
Factors that determine knockout efficiency in single-cell CRISPR screens. **a,** The knockout effect on target gene down-regulation (measured by negative selection p value) and the median target expression in mESC dataset. **b,** the negative selection p values of all targets innaïve and primed states in mESC CROP-seq dataset. **c-d,** The expression of *Tdh* in *Tdh* knockout cells and other cells in naïve and primed states. The CD46 gene itself is marked as red. **e,** The fraction of target genes with the strongest p values, where target gene is ranked top ten among all other genes, in high MOI and low MOI conditions. **f-g,** The p value (f) and selection score (g) of all possible genes upon CD46 knockdown in K562 dataset. CD46 itself is marked as red.

Some CRISPR screening and single-cell CRISPR screening studies suggested using high multiplicity of infection (MOI) to increase the power of screening (Gasperini et al., 2019; Zhu et al., 2019). We set out to compare the effect of high vs. low MOI in terms of target gene knockout effect using scMAGeCK. In the K562 dataset, the screening is performed in two different conditions, one with high MOI (with around 28 gRNAs per cell), and the other with low MOI (around 1 gRNA per cell). We evaluated the statistical power of both conditions, by looking at the effect of down-regulation in over 300 protein-coding genes. The selections scores of these genes are highly correlated between two conditions (Supplementary Fig. 3a). However, over 95% the target genes are among the strongest down-regulated genes in high MOI screen, while only 50%-60% of them ranked top in low MOI screen (Fig. 5e, Supplementary Fig. 3b). For example, CD46 has the strongest down-regulation for CD46 perturbation in high MOI, but only ranks 876th (out of 12,000 genes) in terms of p value, and 305th in terms of selection score in low MOI condition, respectively (Fig. 5f-g). This comparison demonstrates that a better statistical power can be obtained by increasing the level of MOI in the screening experiment.

## Discussion

CRISPR screening using single-cell RNA-seq as readout (“single-cell CRISPR screening”) is a promising technology that overcomes several limitations of traditional CRISPR screening. First, it enables an interrogation of genotypes on potentially unlimited numbers of phenotypes, represented by the expressions of genes or gene signatures. In contrast, CRISPR screening only studies one single phenotype of cell viability or reporter expression. Second, single-cell CRISPR screening reports the effect of perturbations at the single cell level, compared with traditional CRISPR screening that are often performed on bulk cells. To this end, scMAGeCK expands our previous MAGeCK algorithmic framework to analyzing single-cell CRISPR screensing data, providing a powerful computational tool to link genotypes with multiple phenotypes. The two modules of scMAGeCK provide different tools to study gene perturbation in different contexts. scMAGeCK-RRA is an algorithm to detect subtle, non-linear expression changes, while scMAGeCK-LR provides a convenient tool to model the expressions of all genes and is able to deal with cells infected by multiple sgRNAs.

We tested scMAGeCK on several public CROP-seq experiments. scMAGeCK identified potential oncogenes and tumor suppressor genes (and enhancers) by simply testing their associations with the expression of Ki-67, a proliferation marker. We demonstrated the ability of scMAGeCK to study other phenotypes, including apoptosis, T cell stimulation, stem cell differentiation, etc. These results generated from scMAGeCK enabled an unbiased construction of genotype-marker-phenotype network, providing an intuitive picture for users to study gene regulatory network and enhancer-gene regulations.

So far, CRISPR screen studies on mESC pluripotency or naive and primed state transition is mainly based on genetically labeled fluorescence reporters as readout (Hackett et al., 2018; Li et al., 2018; Seruggia et al., 2019), which is limited by only one or two genes. Here we employed a single-cell RNA-seq combined with CRISPR screening technology (CROP-seq) and used whole cell transcriptome as readout of cell fate changes. With the aid of scMAGeCK, we were able to capture alteration of cell fate defined by a combination of marker genes upon genetic perturbation, and to build or refine the regulatory network of mESCs.

MUSIC is a recent tool to model perturbations and their associated biological functions in single-cell CRISPR screening (Duan et al., 2019). MUSIC uses the Topic Model, a method in natural language processing, to connect biological function (“topic”) to gene expression (“word”) in a single cell (“document”) under perturbation. However, MUSIC is not able to efficiently model cells with multiple perturbations, as the MUSIC model relies on the identification of differentially expressed genes between perturbed cells and negative control cells. Also, MUSIC did not accept certain phenotype of interest as input, which may be vary across different users. Both are addressed in our scMAGeCK model, providing a convenient computational framework for biologists.

Some single-cell CRISPR screening technologies (Perturb-seq, CRISP-seq, MOSAIC-seq) use additional barcodes to determine the single cell identity. The sgRNA-barcode correspondence may be compromised during the screening process, which may complicate downstream analysis results(Adamson et al., 2018; Feldman et al., 2018; Xie et al., 2018). Here, we exclusively focus on CROP-seq where sgRNA itself serves as the barcode. Once the sgRNA-barcode issue is solved with improved protocol, scMAGeCK will be extended to other platforms as well.

Besides scRNA-seq, single-cell epigenomic profiling could serve as the screening readout (e.g., single-cell ATAC-seq), providing a novel approach to measure epigenome changes upon perturbation (Rubin et al., 2019). In the future, scMAGeCK will support other types of singlecell technology as the screening readout, enabling analysis on phenotypes beyond gene expressions.

## Methods

### The scMAGeCK algorithm

scMAGeCK consists of two modules, scMAGeCK-RRA and scMAGeCK-LR, based on our previous MAGeCK and MAGeCK-VISPR algorithms (Kolde et al., 2012). scMAGeCK-RRA first ranks single cells based on the expression of gene A of interest. Then, the RRA algorithm proposed by Kolde *et al*. (Hill et al., 2018) to evaluate whether single cells bearing certain gene X is enriched in the front of the ranked list. Suppose *M* single cells are ranked in the experiment, *R* = (*r*_1_, *r*_2_,…, *r_n_*) is the vector of ranks of *n* single cells targeting gene X (*n* < < *M, r_i_* ≤ *M* where *i* = 1, 2,…, *n*), and α is the percentage of single-cells that have non-zero counts on gene A. We first normalize the ranks into percentiles *U* = (*u*_1_, *u*_2_,…, *u_n_*), where *u_i_* = *r_i_*/*M*(*i* = 1, 2,…, *n*). Under null hypotheses where the percentiles follow a uniform distribution between 0 and 1, the *k*th smallest value among *u*_1_, *u*_2_,…, *u_n_* is an order-statistic which follows a beta distribution *B*(*k, n*, + 1 − *k*). RRA computes a *P*-value ρ_*k*_ for the *k*th smallest value based on the beta distribution. For positive selection (cells enriched in higher expression of gene A), the significance score of the gene, the ρ value, is defined as ρ = min(*p*_1_, *p*_2_,…, *p_j_*), where *j* out of the *n* single-cells targeting gene X have non-zero read count on gene A. For negative selection, single-cells ranked in the front will have zero counts (dropouts). Therefore, we remove single cells with dropout events, and only calculate ρ = min(*p*_j+1_, *p*_j+2_,…, *p_n_*) where the first 1-*j* single cells have zero counts on gene A.

To compute a *P*-value based on the ρ values, we performed a permutation test where the sgRNAs are randomly assigned to single cells. We then compute the FDR from the empirical permutation *P*-values using the Benjamini-Hochberg procedure.

The selection score of gene X perturbation on gene A, calculated from scMAGeCK-RRA, combines the results of both negative and positive selection:

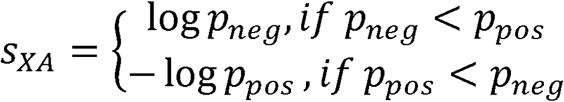

Where *p_neg_* and *p_pos_* are the p values of negative selection and positive selection of perturbing gene X on gene A expression, respectively.

scMAGeCK-LR uses a linear regression model to calculate the selection scores of all genes. Let *Y* be the *M*\**N* expression matrix of *M* single cells and *N* genes. Let *D* be the *M*\**K* binary cell identity matrix, where *d_jX_* = 1 if single cell *j* contains sgRNAs targeting gene *X*(*j* = 1,2,…, *M*; *X* = 1,2,…, *K*), and *d_jX_* = 0 otherwise. The effect of target gene knockout on all expressed genes is indicated in a selection score matrix *S* with size *K*\**N*, where *s_XA_* > 0 (< 0) indicates gene *X* is positively (or negatively) selected on gene *A* expression, respectively. In other words, gene *X* knockout increases (or decreases) gene *A* expression if *s_XA_* > 0 (< 0), respectively.

The expression matrix *Y* is modeled as follows:

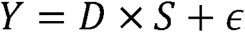

where *ϵ* is a noise term following a Gaussian distribution with zero means. The value of *S* can be estimated using ridge regression:

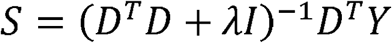

where *I* is the identity matrix, and *λ* is a small positive value (default 0.01).

To compute the empirical *P*-value, we performed a permutation test similar with scMAGeCK-RRA, where the sgRNAs are randomly assigned to single cells. The FDR is then calculated using the Benjamini-Hochberg procedure.

### Public CROP-seq datasets

We used three public CROP-seq datasets. The MCF10A CROP-seq dataset (Hill et al., 2018) is downloaded from Gene Expression Omnibus (GEO) under the accession number GSE90546, and from the corresponding website ((Shifrut et al., 2018)). The T-Cell CROP-seq dataset (Butler et al., 2018) is downloaded from GEO under the accession number GSE119450. The K562 CROP-seq dataset is downloaded from GEO under the accession number GSE120861. All datasets are profiled through 10X Genomics platform. Raw expression matrix from cellranger pipeline is imported and processed using Seurat pipeline (version 3.0) (Butler et al., 2018). Briefly, single cells are first filtered out if they contain <500 expressed genes or >10% read counts coming from mitochondria genes. The expressions of the remaining cells are normalized and scaled based on the number of UMIs and mitochondrial gene expressions. The Principal Component Analysis (PCA), clustering analysis, and t-SNE visualization are performed using default Seurat parameters.

### gRNA library construction

gRNA cassettes were ligated to CROPseq-guide-puro vector using Gibson assembly with a ratio of 20:1 at 50°C for 1h, then dialyze the reaction against water. Electroporate the gRNA library to lucigen endura cells((Lucigen cat. no. 60242-2) using Lonza 2B nucleofector bacteria program 3. After transformation, add 1ml pre-warmed Recovery Medium (Lucigen) and at 37°C for 1h while shaking at 225 rpm. Then 1ml bacterial solution was plated on 25cm×25cm ampicillin LB-agar dish at 34°C for 18h, then LB medium was added to collect the bacteria. Plasmid DNA was extracted with Tiangen EndoFree maxi Plasmid extraction kit (Tiangen cat. no. DP117).

### Lentivirus production for pooled CROPseq screens

HEK293T cells were plated onto 10cm dishes at 6 million cells per dish in 10 ml of lentivirus packaging medium (Opti-MEM I (Gibco), 5% FBS (Gibco), 200mM sodium pyruvate (Gibco). Next day, HEK293T were transfected 11.7 *μ* g constructed CROPseq-guide-puro(containing gRNA library) with lipofectamine 3000 (Invitrogen) using two packaging plasmids psPAX2(addgene 12260) and pMD2.G (addgene 12259). Change the medium to lentivirus packaging medium 6h after transfection. Viral containing supernatant were collected at 24h and 48h. virus were filtered through 0.22 *μ* m filter and 10% PEG 6000 were added to concentrate CROPseq virus. Then CROPseq virus were placed at 4°C overnight. Centrifuging 30min at 4,200 rpm, discard supernatant, resuspend CROPseq virus with 500μl PBS.

### Cell culture

Naive mouse ESCs were cultured in 2i/LIF medium (1:1 DMEM/F12 (Gibco) and Neurobasal medium (Gibco) containing 1%(v/v) N2 and B27 supplements (Gibco), 1mM PD03259010 (stem cell), 3mM CHIR99021 (stem cell), 1000u/ml mLIF (Peprotech), 1X L-glutamine (Gibco), 100mM 2-mercaptoethanol (Sigma) and 1% Penicillin Streptomycin (Gibco)) on 0.1% gelatin-coated dishes with MEF feeders. After transducingCROP-seq gRNA library, ESCs were transferred to FGF2/Activin DMEM/FBS medium (1:1 DMEM/F12 and Neurobasal medium containing 1%(v/v) N2 and B27 supplements, 10ng/ml FGF-2 (Peprotech), 20ng/ml Activin A (Peprotech), 1X L-glutamine, 100mM 2-mercaptoethanol and 1% Penicillin-Streptomycin) for 48h to become primed state cells.

### Single cell RNA-seq

Single cell RNA sequencing was performed with 1cell-bio inDrop platform(Klein, Mazutis et al. 2015). In brief, cells were prepared in 1×PBS containing 1% volume/volume FBS with an input concentration of 40-60 cells/μl. ~6000 cells were captured per sample with different microdevice flow rate conditions at BHM phase varies from 40 to 60 μl/h. Photo-cleavable barcoding oligos were released from barcoded hydrogel microspheres (BHMs) with exposed the collected droplets to UV (6.5J/cm2 at 365 nm). Library preparation was carried out with *in vitro* transcription (IVT), followed by 1st PCR amplification with the following program before fragmentation. 1 cycle of 98°C for 1 min, 10 cycles of 98°C for 7 sec, 60°C for 30 sec, 72°C for 90 sec and 1cycle of 72°C for 3 min. 2st PCR was conducted for final library amplification with following program. 1 cycle of 98°C for 2 min, 2 cycles of 98°C for 20 sec, 55°C for 30 sec, 72°C for 2 min, 9 cycles of 98°C for 20 sec, 65°C for 30 sec, 72°C for 2 min and 1cycle of 72°C for 5 min. One lane was used for sequencing both two samples on Hiseq X.

## Availability

The scMAGeCK source code is freely available at https://bitbucket.org/weililab/scmageck under the BSD license.

## Acknowledgement

We would like to thank Molly Gasperini, Andrew Hill, Cole Trapnell and Jay Shendure for the contribution of scMAGeCK source code, and the discussion of the results. W.L. is supported by the research starter award from Pharmaceutical Research and Manufacturers of American Foundation (PhRMA) and the startup fund from the Center of Genetic Medicine Research and Gilbert Family NF1 Institute at the Children’s National Medical Center. J.Z. is supported by the National Key Research and Development Program of China (2018YFA0107100, 2018YFA0107103, 2018YFC1005002), the National Natural Science Foundation projects of China (31871453, 91857116) and the Zhejiang Natural Science Foundation projects of China (LR19C120001).

